# Dimensionality Reduction for Cyclic Compositional Data

**DOI:** 10.64898/2026.10.22.761053

**Authors:** Nikhila P. Swarna, Jawhara Emhemed, Lior Pachter

## Abstract

The cyclic nature of many biological systems is frequently reflected in periodic gene expression patterns. Standard dimensionality reductions can reveal such variation but do not link a sample’s position on a cycle to an ordering of the molecular features that define it. We introduce *clrcycle*, which applies a centered log-ratio transformation, learns a cyclic feature ordering by maximizing variance captured by the first Fourier mode, and projects samples onto the corresponding cosine and sine coordinates. The resulting two-dimensional geometry assigns each sample a phase and a radius measuring the magnitude of its projection onto the learned Fourier circle. Across a mouse circadian atlas, clrcycle recovers known temporal organization and reveals tissue-specific differences in rhythmic timing. In FUCCI-profiled single cells, clrcycle represents independently measured cell-cycle phase in two-dimensional coordinates more reliably than conventional two-dimensional embeddings. In human endometrium, a circular coordinate learned from 250 highly variable genes recovers menstrual-stage organization and can stage unseen endometrial biopsies. Together, these results establish clrcycle as powerful, explicitly circular dimensionality reduction technique to quantify high-dimensional cyclic variation across biological cycles.

## 1. Introduction

Dimensionality reduction is often the first step in the interpretation of high-dimensional transcriptomic data. Classical linear methods such as singular value decomposition and principal component analysis have long been used to summarize large expression matrices (Alter et al., 2000). More recent nonlinear methods, including diffusion maps (Coifman et al., 2005), Laplacian eigenmaps (Belkin and Niyogi, 2003), UMAP (McInnes et al., 2018), and PHATE (Moon et al., 2019), are some popular tools for displaying cell states, developmental continua, and branching processes. In single-cell analyses, dimensionality reduction is also closely tied to pseudotime inference. Methods such as Monocle (Trapnell et al., 2014), reversed graph embedding (Qiu et al., 2017), and Slingshot (Street et al., 2018) infer ordered or branched trajectories and use low-dimensional displays to visualize progression.

However, many biological trajectories are not fundamentally linear. Circadian time, the cell cycle and mammalian menstrual and estrous cycles are inherently circular. Some perturbation and regeneration experiments similarly trace trajectories that return toward a starting state. These settings call for dimensionality reduction and visualizations in which the primary latent coordinate is circular. Several approaches have been developed to represent such structure. Principal-curve methods can be extended to fit a closed principal circle through high-dimensional data, providing a continuous cyclic trajectory onto which observations can be projected (Albergante et al., 2020). Topological methods instead detect loops and construct circular coordinates from persistent cohomology (Silva et al., 2011), allowing cyclic structure to be identified without specifying a parametric trajectory. These approaches are complementary to harmonic analyses of rhythmic transcription, in which periodic variation is modeled directly using sine and cosine components (Hughes et al., 2009). However, these methods do not simultaneously learn an interpretable circular ordering of molecular features and use that ordering to define circular coordinates for the samples themselves.

We introduce *clrcycle*, a method that asks a different but related question (Fig. 1). Instead of first finding a circular coordinate for samples, we ask whether the molecular features or genes themselves can be ordered on a circle so that the dominant covariance looks like a first Fourier mode. We first order selected genes around a circle by the phase at which their relative abundance contributes to the cyclic signal. Samples can then be projected onto the same circular geometry such that their angle corresponds to phase, and radius corresponds to strength of the “cyclic signal” with respect to the selected feature set, fit to the first Fourier mode. This radius can further be decomposed into feature weights quantifying how much each gene is contributing to the first Fourier cyclicity of a given sample. clrcycle’s feature-forward circular geometry provides a uniquely interpretable framework that complements other dimensionality reductions techniques and provides a supervised approach to dimensionality reduction in cases where a cyclic biological coordinate is plausible.

**Figure 1.**
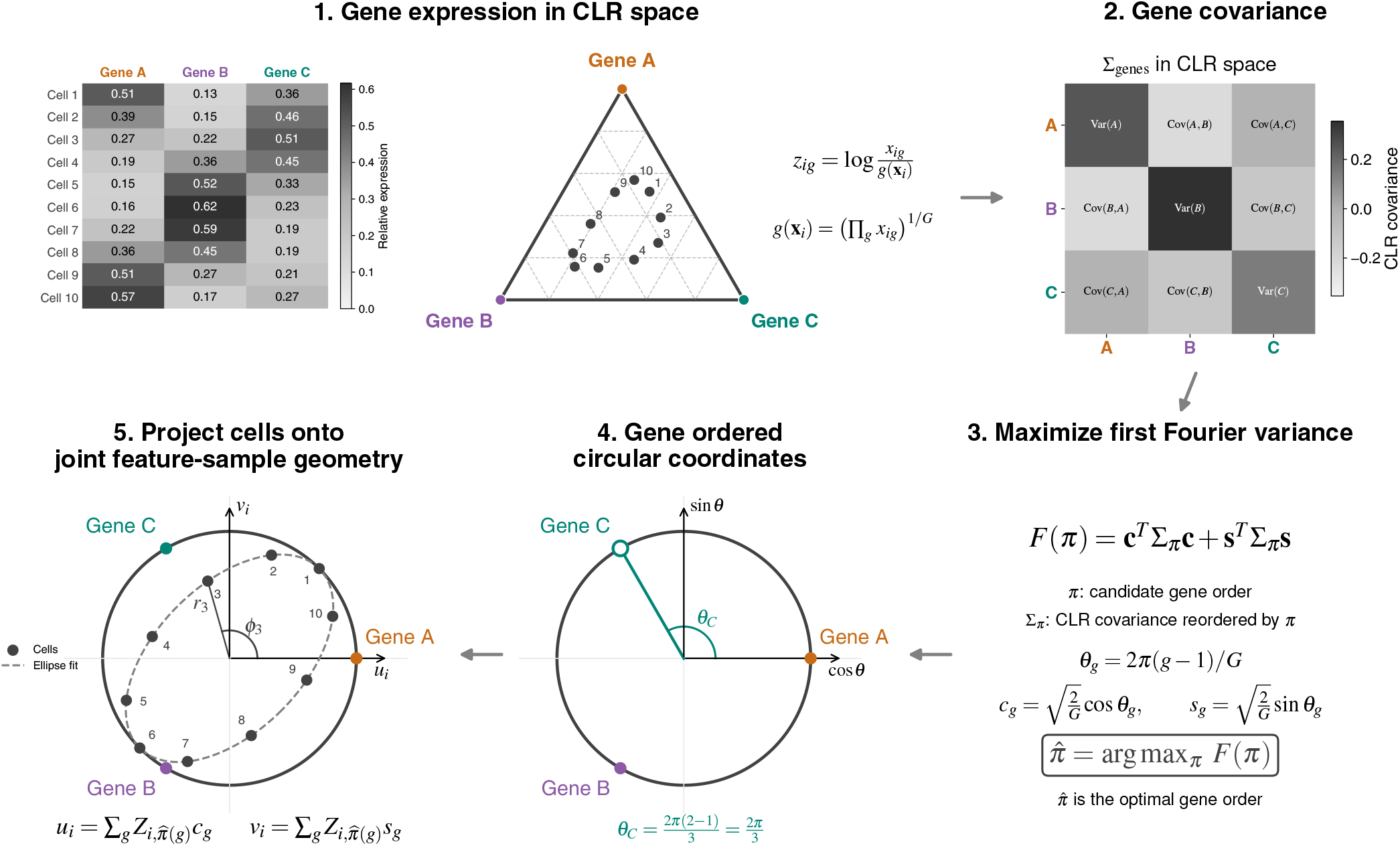
clrcycle architecture. (1) Ten synthetic cells are represented as compositions of three genes. The heatmap shows relative expression fractions, and the triangle displays the corresponding compositions on a 2-simplex. Gene abundances *x*_*ig*_ are transformed to centered log-ratio (CLR) coordinates relative to the geometric mean expression within each cell; each gene is then centered across cells. (2) Gene-gene covariance Σ_genes_ is computed in centered CLR space. (3) clrcycle searches for a circular gene ordering 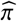 that maximizes the variance captured by the Fourier cosine and sine directions. For this three-gene illustration, all circular orders span the full CLR plane and therefore yield the same objective value. (4) The selected order assigns each gene an equally spaced circular angle *θ*_*g*_ and corresponding Fourier coordinates (*c*_*g*_, *s*_*g*_). (5) Cells are projected onto these coordinates to obtain (*u*_*i*_, *v*_*i*_), from which phase *ϕ*_*i*_ and radius *r*_*i*_ are calculated. Phase represents position along the circular coordinate, whereas radius measures the magnitude of the cell’s projection onto the Fourier plane. The synthetic profiles are constructed to form an ellipse, and the dashed ellipse is fitted using all ten cells. Cells 1 and 6 are constructed to lie on the outer unit reference circle, while the remaining cells have smaller projection radii.

Our approach’s compositional formulation (via the centered-log ratio, hence “clr”) is important because gene expression measurements are inherently compositional after read-depth normalization, and because compositional data are most naturally analyzed using log-ratio coordinates (Aitchison, 1982). CLR centering removes sample-wise scale thereby reducing the influence of sequencing depth on the resulting geometry (Booeshaghi et al., 2026).

This perspective is related to simplex-aware visualizations of compositional gene specificity (Swarna et al., 2026). Compositional profiles naturally occupy a simplex, where relative rather than absolute abundance defines their geometry. Although polygonal projections provide an intuitive way to visualize such compositions, their appearance depends strongly on the ordering of features around a polygon. This dependence on feature order is well known in radial visualization methods, beginning with dimensional anchors (Hoffman et al., 1999; Hu et al., 2021; Zhu et al., 2022; Paciencia et al., 2019). Rather than specifying this ordering arbitrarily, our method learns it directly from the data’s covariance structure.

We evaluate clrcycle across three distinct biological cycles: circadian transcription across mouse tissues (R. Zhang et al., 2014), cell-cycle progression in FUCCI-profiled single cells (Hsiao et al., 2020), and menstrual-cycle progression in human endometrium (Teh et al., 2023; Wang et al., 2020). In each of these case studies, genes from sequencing data are treated as features which are pre-selected to define the circular feature space onto which samples or cells are projected. Through these examples, we showcase clrcycle as a novel geometric framework that recovers known temporal organization of both genes and samples and provides interpretable circular coordinates for a reduced two-dimensional representation of high-dimensional cyclic variation.

## 2. Results

### 2.1. Multi-tissue circadian time can be reduced to 2D-circular coordinates

We first applied clrcycle to the GSE54650 mouse circadian atlas, which profiles gene expression across 12 tissues every 2 h over 48 h (R. Zhang et al., 2014). Using 240 probes with the strongest association with the recorded circadian times, clrcycle organizes liver samples around a circle in their known temporal order (circular agreement *R* = 0.951; Fig. 2A). The radius of each sample corresponds to the sum of its feature-level projection weights, so we can decompose a sample into each gene’s contribution to its first-harmonic projection. In the liver we can identify *Dbp, Ciart* and *Arntl* as some of the top drivers of cyclicity in the first Fourier mode (Supplementary Fig. 1, complete feature weights across tissues in Supplementary Table 1). The learned gene ordering similarly reflects distinct circadian phases, with canonical rhythmic genes including *Dbp, Rorc*, and *Clock* (Takahashi, 2017), occupying different positions around the circle consistent with their expression peaks (Fig. 2B–E).

**Figure 2.**
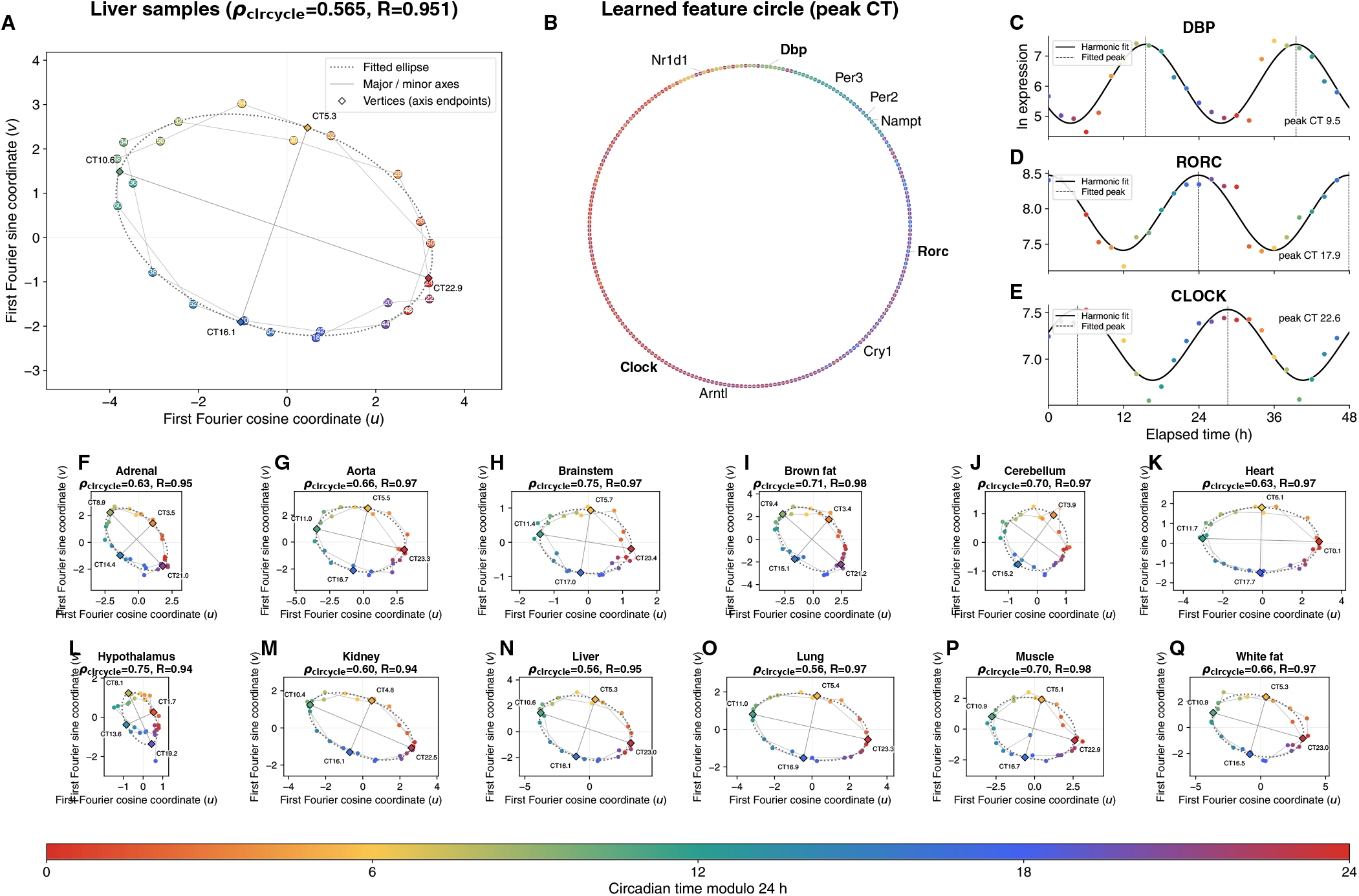
Circadian phase recovery in GSE54650. (A) Liver samples in aligned clrcycle coordinates, colored by circadian time modulo 24 h and labeled by sampling time (CT18–CT66). (B) The 240 selected liver probes on the learned feature circle, colored by harmonic peak time. (C–E) Log expression and first-harmonic fits for *Dbp, Rorc*, and *Clock*; dashed vertical lines mark fitted peaks. (F–Q) Independently fitted projections for adrenal, aorta, brainstem, brown fat, cerebellum, heart, hypothalamus, kidney, liver, lung, muscle, and white fat. In A and F–Q, dotted curves show fitted ellipses, straight gray lines show their major and minor axes, and diamonds mark the four axis endpoints, colored and labeled by their aligned circadian times. Thin gray lines connect samples in sampling order. Projection titles report *ρ*_clrcycle_ and circular agreement *R*.

We next applied the same analysis independently across all 12 tissues. clrcycle recovered circadian sample ordering in all tissues (*R* = 0.94–0.98; Fig. 2F–Q). We noticed an elliptical pattern evident in sample organization across tissues. Qualitatively, an elliptical pattern in this reduced gene space can be interpreted as follows: certain samples are more aligned with the first Fourier mode of the selected genes than others, and these highly aligned samples correspond to opposing circadian times. Quantitatively, we can fit an ellipse directly to the clrcycle sample coordinates (Table 1) and learn statistically significant anisotropy of samples along circadian time. All tissues except cerebellum exhibited significant anisotropy, with axis ratios ranging from *b*/*a* = 0.51–0.71, while cerebellum was nearly circular (*b*/*a* = 0.95), significance test p-values in Supplementary Table 1, stability test described in Methods).

**Table 1.** Summary of circadian recovery across tissues. Here, *a* and *b* are the major and minor semiaxes of the fitted ellipse, and *d* is its center’s distance from the clrcycle origin. Major-axis CT gives the inferred circadian times of the two major-axis vertices, mapped from their angular directions relative to the origin and reported modulo 24 h.

| Tissue | $\rho_{\text{clrcycle}}$ | $R$ | $b/a$ | $d/a$ | Major-axis CT |
| --- | --- | --- | --- | --- | --- |
| Adrenal | 0.63 | 0.95 | 0.61 | 0.10 | 8.9, 21.0 |
| Aorta | 0.66 | 0.97 | 0.71 | 0.09 | 11.0, 23.3 |
| Brainstem | 0.75 | 0.97 | 0.69 | 0.06 | 11.4, 23.4 |
| Brown fat | 0.71 | 0.98 | 0.68 | 0.04 | 9.4, 21.2 |
| Cerebellum | 0.70 | 0.97 | 0.95 | 0.10 | 9.5, 21.8 |
| Heart | 0.63 | 0.97 | 0.56 | 0.06 | 0.1, 11.7 |
| Hypothalamus | 0.75 | 0.94 | 0.54 | 0.12 | 8.1, 19.2 |
| Kidney | 0.60 | 0.94 | 0.51 | 0.05 | 10.4, 22.5 |
| Liver | 0.56 | 0.95 | 0.63 | 0.11 | 10.6, 23.0 |
| Lung | 0.56 | 0.97 | 0.54 | 0.05 | 11.0, 23.3 |
| Muscle | 0.70 | 0.98 | 0.70 | 0.03 | 10.9, 22.9 |
| White fat | 0.66 | 0.97 | 0.64 | 0.07 | 10.9, 23.0 |

The fitted major-axis vertices of the ellipse, where the anisotropy peaks, provide tissue-specific estimates of the phases of greatest Fourier projection magnitude. In most tissues, these peaks occur near CT8–11 and CT20–23. We observed that the heart was the only exception, peaking at CT11.7 and CT0.1, it’s tides seemingly offset from other tissues. Since the mice were trained to a 12:12 light:dark cycle prior to culling, CT0 and CT12 correspond to subjective dawn and dusk respectively. This geometry is consistent with the gene-wise analysis of Zhang et al., who found that circadian-gene expression frequently peaked in bimodal “rush hours” preceding subjective dawn and dusk and that genes peaking near these transitions tended to exhibit greater oscillation amplitudes than genes peaking at other phases (R. Zhang et al., 2014). Both Zhang et al. and a separate study by Storch et al. found that circadian phase distribution of the heart differs significantly from other tissues (Storch et al., 2002; R. Zhang et al., 2014). Previous studies have also observed sexual dimorphism in transcription in the murine heart, noting that the male heart had transcriptional peaks at CT0 and CT12 (X. Zhang et al., 2024). This aligns with our findings from the Zhang et al. data, which was collected in male mice. It has also been noted that light exposure, in addition to feeding patterns affect circadian transcription dynamics and the heart, is specifically resistant to entrainment by feeding. We therefore hypothesize that the male murine heart has distinct transcriptional peaks along circadian time due to its resistance to feeding based entrainment.

Our feature-based coordinate system to reduce the dimensionality of this multi-tissue dataset allows for quantification and visualization of subtle differences across tissues and circadian time, facilitating hypothesis generation and biological discovery. clrcycle provides an intuitive and feature-forward framework for understanding circadian dynamics in more complex experimental designs across experimental modalities and organisms.

### 2.2. Cell-cycle phase in single cells can be represented in an explicitly circular embedding

We also applied clrcycle to 888 human induced pluripotent stem cells from GSE121265 with paired fluorescent ubiquitination cell cycle indicators (FUCCI). FUCCI technology independently measures paired fluorescent reporters corresponding to each cell cycle phase, thereby providing closer to ground-truth continuous cell-cycle phase (Hsiao et al., 2020). Using the 100 genes from this dataset identified as having the strongest cyclic expression trends, clrcycle organizes individual cells according to measured FUCCI phase (*R* = 0.441; Fig. 3A,B). The learned gene cycle similarly orders genes by their independently fitted peak phases, with *DIABLO, CDC6*, and *TNPO1* illustrating distinct expression peaks across the cell cycle (Fig. 3C–E). The ellipse fitted to samples projected into the circular feature space was strongly anisotropic (*b*/*a* = 0.50), with major-axis vertices mapping to G1-and G2-associated phases. We can interpret this as G1- and G2-associated phases having greater transcriptional cyclicity in this this 100-feature space. This is consistent with the broader organization of cell-cycle transcription into distinct phase-dependent programs (Hsiao et al., 2020). Due to the noise which is inherent to single cell sequencing technologies, mapping single cells along cyclic time is noisier than working with bulk samples. This can result in lower confidence estimates, such as seen with circular agreement of the cells with FUCCI phase (*R* = 0.441). Despite noise, we can still learn relative patterns of cyclicity across samples of cells, as with the observed anisotropy of this data being is strongly significant with a leave-one-out p-value of 0.0009 (Supplementary Table 2).

**Figure 3.**
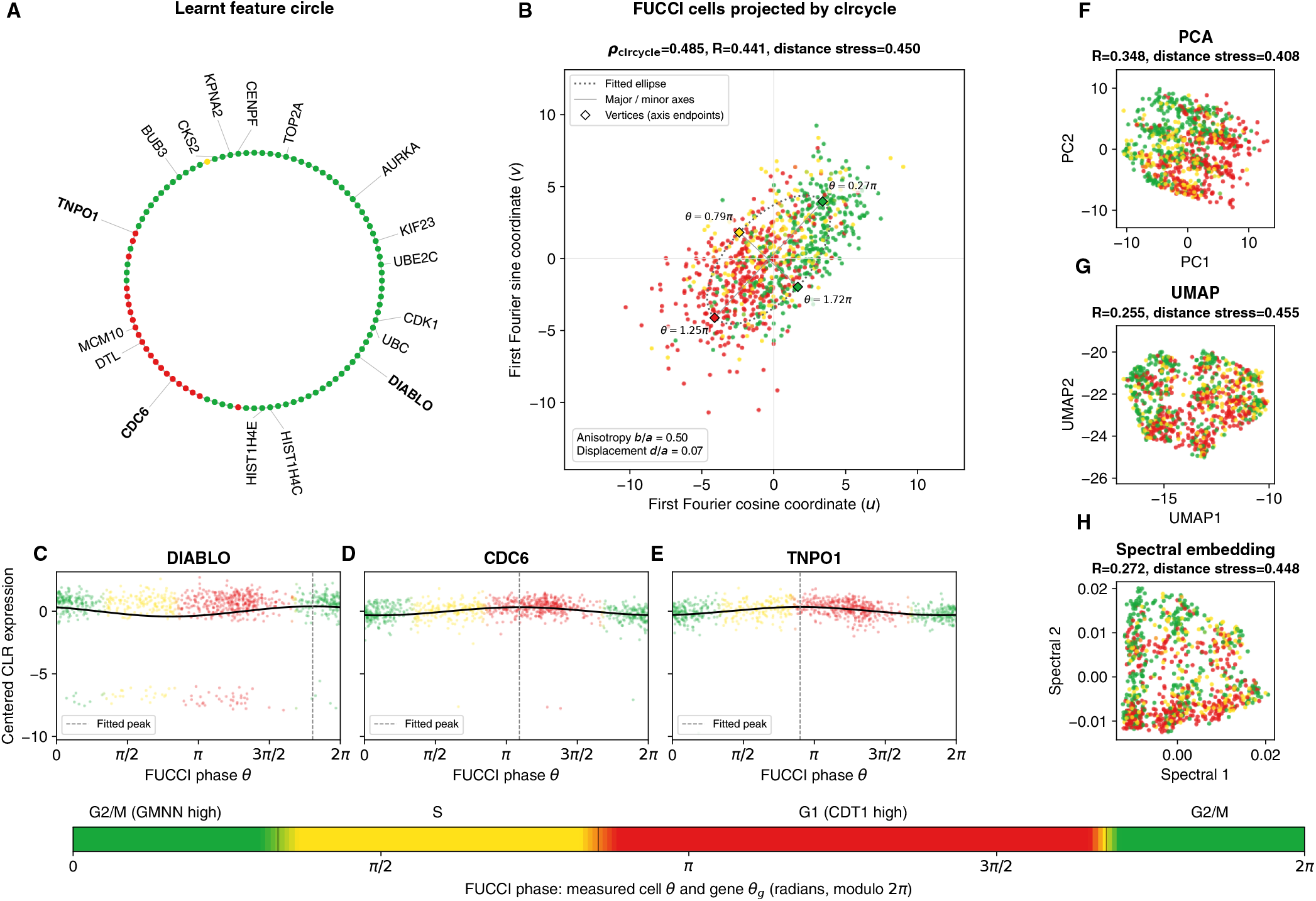
FUCCI phase analysis and embedding benchmark in GSE121265. (A) Learned order of the published top-100 cyclic genes, colored by their published fitted peak phases. (B) clrcycle projection of 888 cells, colored by measured FUCCI phase. The dotted curve shows the fitted ellipse, straight gray lines show its major and minor axes, and diamonds mark the four axis endpoints, colored and labeled by their aligned FUCCI phases in radians. The inset reports anisotropy *b*/*a* and displacement *d*/*a*, where *a* and *b* are the major and minor semiaxes and *d* is the ellipse center’s distance from the origin. (C–E) Centered CLR expression and first-harmonic fits for *DIABLO, CDC6*, and *TNPO1*; dashed vertical lines mark fitted peaks. (F–H) PCA, UMAP, and spectral embeddings of the same centered CLR matrix. Embedding panel headings report circular agreement with FUCCI phase and normalized pairwise-distance stress; B also reports *ρ*_clrcycle_.

One of the key contributions of this feature-specific circular representation is preservation of cyclic information when embedding high-dimensional samples in two-dimensions. Using the same 888 cells and 100-gene CLR representation, clrcycle showed greater agreement with measured FUCCI phase (*R* = 0.441) than PCA (*R* = 0.348), spectral embedding (*R* = 0.272), or UMAP (*R* = 0.255; Fig. 3B,F–H). While clrcycle best preserves circular agreement, it should be noted that PCA best preserved global pairwise distances (stress = 0.408 versus 0.450 for clrcycle). Thus, clrcycle provides a dimensionality reduction framework that facilitates representation of cyclic structure of samples given selected features and can be used to interpret highly noisy, high-dimensional biological cycles.

### 2.3. Menstrual-cycle stage recovered from circular coordinates of highly variable endometrial genes can stage unseen endometrial samples

Despite its importance in human fertility and regenerative biology, the cyclic remodeling of the endometrium remains understudied (Lin et al., 2026). Menstrual-cycle stage is a major source of transcriptomic variation in endometrial biopsies and can confound the identification of other biological signals (Devesa-Peiro et al., 2021; Teh et al., 2023). Teh et al. profiled endometrial biopsies across seven histological stages and showed that their transcriptomes trace a continuous cycle, motivating the need for molecular approaches to accurately stage endometrial samples (Teh et al., 2023). We therefore asked whether clrcycle could recover this latent cyclic structure directly from endometrial gene expression.

Using only the 250 most variable genes selected from the training RNA-seq samples, clrcycle organized biopsies around a circular coordinate that closely followed histological stage (*R* = 0.867; Fig. 4A,B). This coordinate is also concordant with the continuous molecular cycle time inferred by Teh et al. by fitting cyclic splines to 20,067 genes (*R* = 0.922; Fig. 4B,C). We can quantify sample anisotropy along the menstrual cycle by fitting the samples in circular coordinates to an ellipse. The major axis vertices align with the secretory stage and the late proliferative stage. The fitted ellipse has a significant displacement of 0.16 in the direction of the secretory stage. We can interpret this as samples in the secretory stage aligning more strongly with the first Fourier harmonic than other menstrual stages. This geometry of anisotropy is in agreement with prior observations that the most dramatic transcriptional changes in the human endometrium occur during the secretory phase (Teh et al., 2023).

**Figure 4.**
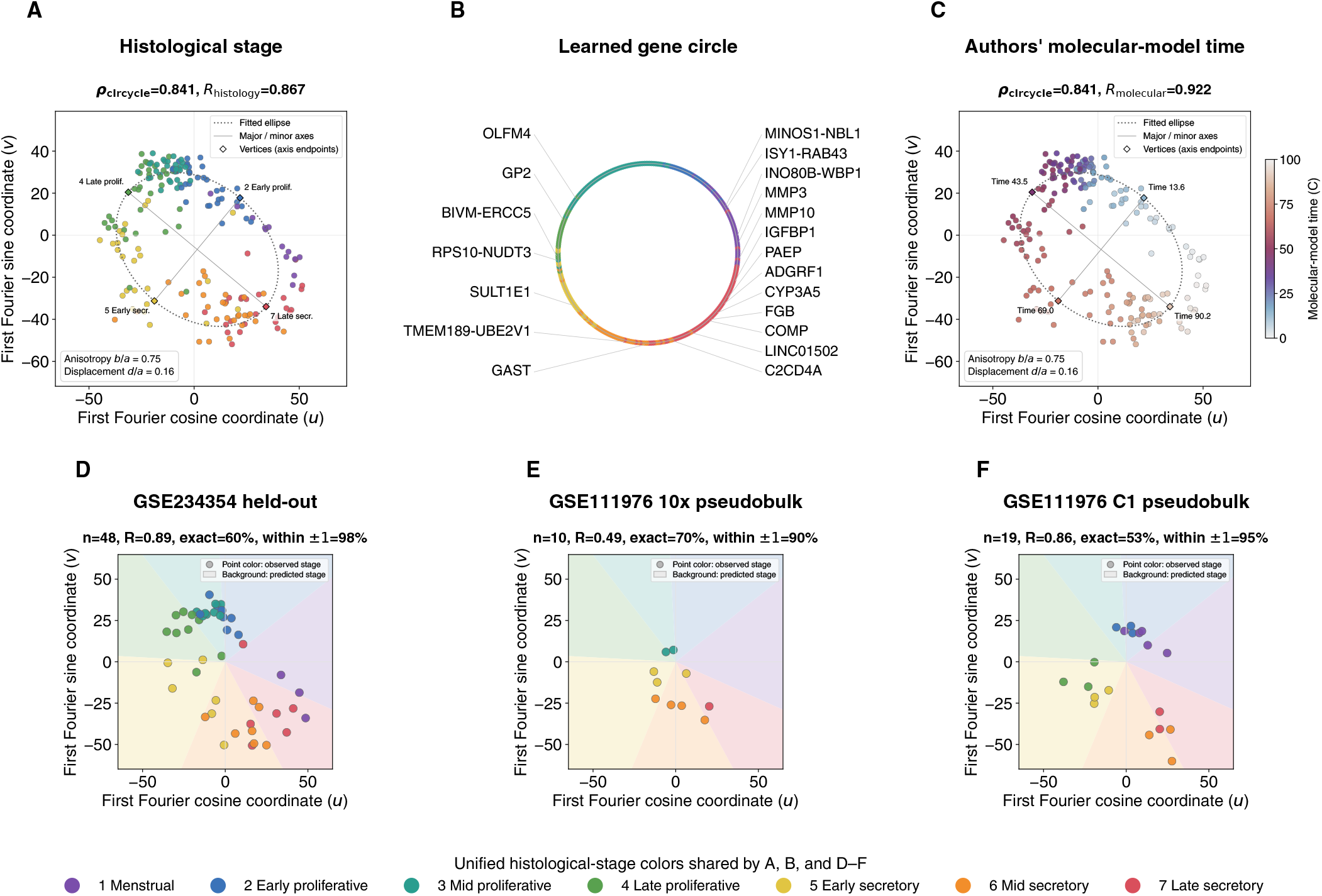
clrcycle captures menstrual-cycle organization and enables staging across datasets and platforms. (A) GSE234354 training endometrial biopsies projected by clrcycle and colored by histological menstrual stage. (B) Learned circular ordering of the 250 highly variable genes, colored by their post-hoc peak histological stage. (C) The same biopsy projection colored by the independently inferred molecular cycle time from Teh et al. In A and C, dotted curves show fitted ellipses, straight gray lines show their major and minor axes, and diamonds mark the four axis endpoints, colored and labeled by aligned histological stage or molecular cycle time. Insets report anisotropy *b*/*a* and displacement *d*/*a*, where *a* and *b* are the major and minor semiaxes and *d* is the ellipse center’s distance from the origin. Headings report *ρ*_clrcycle_ and circular agreement *R*. (D) Menstrual-stage prediction for held-out GSE234354 biopsies projected onto the learned clrcycle coordinates. (E,F) Transfer of the same learned coordinates to GSE111976 10x and C1 donor pseudobulks without refitting. In D–F, points are colored by observed menstrual stage and background sectors indicate predicted stages; headings report sample size, circular agreement *R*, exact stage accuracy, and accuracy within one stage.

Teh et al. addressed the challenge of molecular staging by fitting cyclic expression models across 20,067 genes. We therefore asked whether the substantially lower-dimensional clrcycle representation could provide a useful coordinate for staging unseen samples. Projection of 48 held-out biopsies through the fixed 250-gene clrcycle coordinate aligned with menstrual stage circular ordering (*R* = 0.887) and assigned 60.4% of samples to their exact histological stage and 97.9% to within one stage (Fig. 4D). Thus, the circular coordinate learned from a small highly variable gene panel was sufficient not only to visualize menstrual progression, but also to recover stage in samples excluded from fitting.

This coordinate can also stage unseen samples in data collected from other sequencing technologies and processed independently. We projected donor-level pseudobulks from the GSE111976 endometrial single-cell study onto the fixed circular clrcycle coordinate learnt from the Teh et al. data (Wang et al., 2020). Menstrual organization remained apparent in both 10x and C1 data (Fig. 4E,F): stage assignment was within one stage for 90.0% of 10x donors and 94.7% of C1 donors, with circular agreement with reported menstrual stage circular ordering *R* = 0.494 and *R* = 0.859, respectively. This suggests that clrcycle’s reduced two-dimensional feature-specific coordinates not only fairly represent cyclic menstrual signal, but also provide a uniquely interpretable framework to harmonize information across datasets along the same biological cycle.

## 3. Discussion

clrcycle links two views of the same cyclic transcriptomic program: a circular ordering of molecular features and a phase-amplitude projection of samples. The angles in these representations have a specific interpretation. Genes placed near one another on the feature circle have similar phase with respect to the learned compositional program, while samples with similar angles share similar cyclic transcriptional profiles within the selected feature space. This similarity is therefore conditional on the genes used to define the circular coordinates and may change when the feature set changes.

The radius has an equally specific interpretation. For a sample, *r*_*i*_ is the magnitude of its projection onto the Fourier plane defined by the learned gene ordering. A larger radius therefore indicates stronger alignment with this fitted Fourier representation; it should not be interpreted as total expression, transcriptional activity, or a general measure of biological signal strength. CLR centering removes sample-wise scale (Booeshaghi et al., 2026), so this radial coordinate reflects the magnitude of the cyclic compositional contrast rather than absolute expression magnitude. Both phase and radius are conditional on the supplied feature space, making clrcycle a targeted dimensionality-reduction method for feature sets containing a cyclic signal of interest.

Our benchmarks examine the consequences of imposing such a representation. The learned clrcycle gene ordering agreed more closely with histological menstrual stage than an ElPiGraph principal-circle ordering (Albergante et al., 2020), indicating that the feature circle is not trivially recovered by fitting a generic closed trajectory in the original feature geometry and supporting an optimization framework for this task. Restricting the representation to the first Fourier mode also retained most of the structure accessible to higher-order harmonics and a periodic spline, while capturing substantially more than a linear model. At the sample level, the FUCCI comparison illustrated a complementary tradeoff: clrcycle better represented the independently measured circular phase, whereas PCA better preserved global pairwise distances. Thus, the circular constraint prioritizes a particular cyclic coordinate in a two dimensional representation of high dimensional data rather than attempting to preserve all aspects of the data’s high-dimensional geometry.

Feature selection thus determines which cyclic program clrcycle can represent. In the endometrial analysis, the 250 most variable genes were sufficient to recover menstrual time without having to selecting genes using histological stage. The unsupervised inference of menstrual time is only plausible in this analysis because menstrual stage is a major source of transcriptional variation in the human endometrium (Teh et al., 2023; Devesa-Peiro et al., 2021). Highly variable genes may not necessarily isolate the relevant cyclic program in other settings, especially in single-cell data that is inherently noisier.

Unsupervised identification of cyclic structure from transcriptomic data has been approached in several ways, including oscillatory gene detection and cyclic cell ordering in Oscope, cell-cycle ordering in reCAT, latent periodic trajectory inference in Cyclum, and circadian ordering in CYCLOPS and Tempo (Leng et al., 2015; Liu et al., 2017; Liang et al., 2020; Anafi et al., 2017; Auerbach et al., 2022). Tricycle projects cells into a fixed circular PCA reference to estimate cell-cycle position (Zheng et al., 2022). The elegant CocycleHunter framework of Maggs et al. (2025) uses persistent cohomology to detect circular structure in gene-set projections, separate cyclic processes, and estimate gene phases through lead–lag analysis. Grande and Meilă (2026) provide a complementary way to determine which candidate circular variables explain detected topological loops. In contrast, clrcycle learns a Fourier ordering from the covariance of a selected gene set and uses that ordering to project new samples. Extending clrcycle toward identifying cyclic feature sets without a known temporal coordinate represents an interesting direction for future work.

clrcycle provides a deliberately targeted dimensionality reduction framework: given a feature space containing cyclic structure, it learns an interpretable circular organization of those features and uses the same geometry to quantify the corresponding organization of samples. In this way, clrcycle reduces high-dimensional cyclic variation to an explicitly circular two-dimensional representation while preserving a direct and interpretable connection between sample coordinates and the molecular features that define them.

## 4. Methods

### 4.1. clrcycle architecture

Figure 1 summarizes clrcycle in five steps. Let 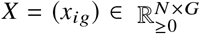, where *i* ∈ {1, …, *N*} indexes samples or cells, *g* ∈ {1, …, *G*} indexes selected genes, and *x*_*ig*_ is the nonnegative abundance of gene *g* in sample *i*.

#### 4.1.1 Gene expression in CLR space

The row vector **x**_*i*_ = (*x*_*i*1_, …, *x*_*iG*_) is treated as a composition, so only relative abundance within sample *i* is used. Zeros are handled with a small pseudocount prior to transformation. Let

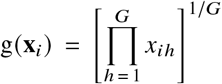

be the geometric mean within sample *i*, where *h* is a dummy gene index. The centered log-ratio coordinate is

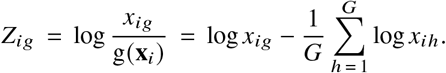

Each gene column is then centered across samples by replacing *Z*_*ig*_ with 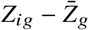, where 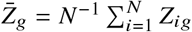. Henceforth, *Z* denotes this column-centered CLR matrix.

Because CLR coordinates represent log abundance relative to the within-sample geometric mean, positive and negative values indicate relative enrichment and depletion, respectively; zero or low-count features can produce strongly negative values following pseudocount replacement, resulting in the negative bands visible in the CLR expression-versus-phase plots (Figs. 2C–E).

#### 4.1.2 Gene covariance

The gene covariance matrix in centered CLR space is

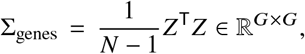

where ^T^ denotes transpose. Its (*g, h*) entry is

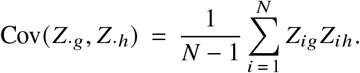

Thus each diagonal entry is Var(*Z*_·*g*_), and each off-diagonal entry is the covariance between two genes.

#### 4.1.3 Maximize first Fourier variance

A candidate order *π* is a permutation of the *G* genes. The *G* positions are equally spaced around a circle as such:

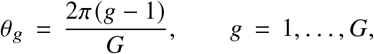

The normalized Fourier vectors **c, s** ∈ ℝ^*G*^ are orthonormal, with each vector summing to zero, and have components

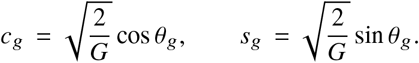

Because 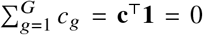 and 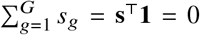, both Fourier basis vectors are orthogonal to the CLR constraint vector **1**, which ensures the projection lies withing the (*G* − 1)- dimensional Aitchison simplex space. For a candidate ordering *π*, let Σ_*π*_ denote the simultaneous permutation of the rows and columns of the gene–gene covariance matrix, rearranged to match that order. No covariance values are recomputed; only their positions in the matrix change.

And the scalar

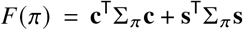

is the total centered-CLR variance captured by these two Fourier directions. Expanding this gives 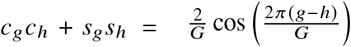 so we can write *F*(*π*) = tr(Σ_*π*_ *K*), where 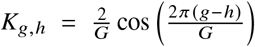 is a symmetric circulant matrix. This establishes that the Fourier variance is an instance of circular seriation formulated as a Quadratic Assignment Problem(Recanati et al., 2018). The learned order is

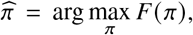

where arg max denotes the gene order that maximizes the objective. The implementation initializes *π* from the polar angles of the two leading covariance eigenvectors and then retains strictly improving pairwise swaps. Reported fits used 4,000 proposals. It should be noted that this is a heuristic greedy search that does not guarantee the global maximum.

#### 4.1.4 Gene ordered circular coordinates

Under 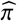, each gene is assigned to one of the *G* equally spaced positions on the circle. Gene 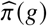 occupies position *g* and is assigned angle *θ*_*g*_. Its coordinates on the unit circle are

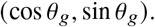

For example, in the three-gene schematic, Gene C occupies position *g* = 2, giving

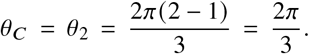

#### 4.1.5 Project cells onto joint feature-sample geometry

Each sample is projected onto the cosine and sine coordinates of the learned gene order:

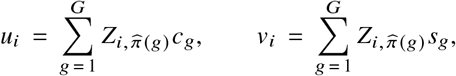

The sample phase and radius are

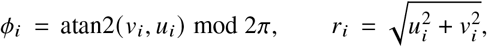

where atan2 is the quadrant-aware polar angle and the modulo operation maps it to [0, 2*π*). Thus, *ϕ*_*i*_ gives the sample’s position along the learned cyclic coordinate, while *r*_*i*_ measures the magnitude of its projection onto the Fourier plane. A global rotation or reflection of the circle is not identified by the objective and may be fixed afterward using a reference phase.

#### 4.1.6 Decomposing sample projections into feature weights

The radial projection can additionally be decomposed into feature-level contributions. For sample *i*, the contribution of gene *g* along the sample’s projected direction is

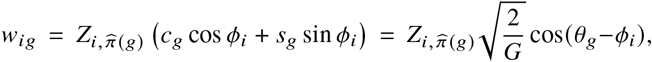

where *θ*_*g*_ is the angular coordinate of position *g*, occupied by gene 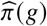, in the learned Fourier basis. These feature-level projection weights sum to the sample radius, because *u*_*i*_ = *r*_*i*_ cos *ϕ*_*i*_ and *v*_*i*_ = *r*_*i*_ sin *ϕ*_*i*_ taking the sum gives 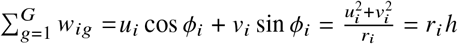

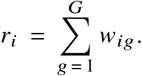

Thus, *w*_*ig*_ quantifies the contribution of feature *g* to sample *i*’s radial position in the learned cyclic coordinate system: positive weights reinforce the sample’s projected direction, whereas negative weights oppose it.

### 4.2. Evaluation

#### 4.2.1 Fourier variance

Let *λ*_1_ ≥ *λ*_2_ be the leading eigenvalues of Σ_genes_. We report

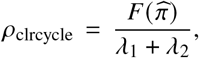

the Fourier variance relative to the maximum variance attainable by an unconstrained two-dimensional linear projection. Thus, *ρ*_clrcycle_ measures how much of the leading two-dimensional covariance structure is captured by the Fourier plane associated with the learned circular feature ordering.

#### 4.2.2 Circular agreement with reference phase

When an external reference phase *ψ*_*i*_ was available, we compared it with the inferred clrcycle phase *ϕ*_*i*_ after accounting for the arbitrary orientation of the learned circle. We selected the reflection *q* ∈ {−1, 1} that maximized

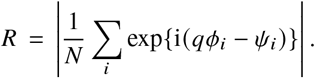

For the selected reflection, we set the rotation to

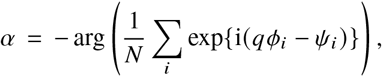

which centers the mean phase difference at zero. Ties in *R* were resolved by smaller mean absolute circular error. Here, *R* = 1 indicates perfect circular agreement after global rotation and reflection. Mean absolute circular error (MACE) was calculated from the absolute wrapped residuals after this alignment and was rescaled to hours, stages, or the source time coordinate where stated. Reference phase was used only for post hoc evaluation and did not enter the clrcycle fit.

#### 4.2.3 Ellipse geometry of sample projections

To quantify phase-dependent variation in the magnitude of the clrcycle projection, we fit an ellipse directly to the raw sample coordinates (*u*_*i*_, *v*_*i*_). We represented the ellipse by the general conic

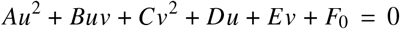

and estimated its coefficients by direct constrained algebraic least squares using the Halíř–Flusser formulation (Halíř and Flusser, 1998). Specifically, the coefficients were chosen to minimize the summed squared algebraic residuals

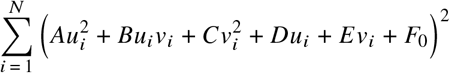

subject to

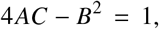

which constrains the fitted conic to be an ellipse and fixes the otherwise arbitrary scale of its coefficients. All samples contributed equally to the fit, and no samples were removed as outliers.

The fitted conic was converted to its geometric representation, yielding center (*u*_0_, *v*_0_), major and minor semiaxes *a* ≥ *b*, and major-axis orientation *η*. We summarize anisotropy by the axis ratio

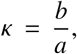

We additionally report the displacement of the ellipse center from the clrcycle origin,

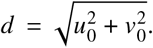

Ellipse fits can be interpeted as follows:

- **Anisotropy:** where *κ* = 1 corresponds to a circle and smaller values indicate increasing anisotropy of the Fourier projection magnitude across phase.
- **Ellipse vertices:** where the four vertices correspond to the endpoints of the major and minor axes, with their angles measured relative to the clrcycle origin.
- **Ellipse displacement:** where displacement of the ellipse center indicates greater Fourier projection magnitude toward one phase than its opposing phase. Because the absolute scale of the clrcycle coordinates can differ between datasets, center displacement is summarized relative to the major semiaxis as *d*/*a*.

The ellipse fit minimizes algebraic rather than orthogonal Euclidean distance to the ellipse. We therefore additionally report the root-mean-square dimensionless implicit residual

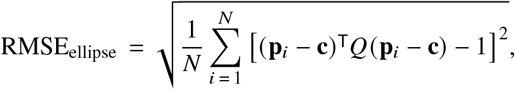

where **p**_*i*_ = (*u*_*i*_, *v*_*i*_)^T^, **c** = (*u*_0_, *v*_0_)^T^, and *Q* is the quadratic form of the fitted ellipse with eigenvalues 1/*a*^2^ and 1/*b*^2^.

#### 4.2.4 Ellipse stability and anisotropy

We performed two additional analyses to assess the robustness and statistical significance of the fitted ellipse.

##### Leave-one-out stability

To determine whether the ellipse geometry was driven by individual samples, we repeated the ellipse fit after removing each sample in turn. Each leave-one-out fit was compared with the full-data fit using semiaxis stability, orientation stability, vertex stability.

These quantities measure sensitivity to individual observations rather than statistical significance. Orientation and vertex angles are expected to become less stable as *κ* = *b*/*a* approaches one, because a nearly circular ellipse has no strongly defined major axis.

##### Anisotropy permutation test

To test whether the observed ellipse was more anisotropic than expected under circular angular geometry, we defined

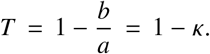

Thus, *T* = 0 corresponds to a circle, while larger values indicate greater anisotropy.

For the null distribution, each sample’s observed radius *r*_*i*_ was preserved while its angle was independently randomized over the circle. An ellipse was then refit to each randomized dataset using the same fitting procedure. This preserves the observed distribution of projection magnitudes while removing their association with angular position.

For *M* permutations, the empirical *P*-value was

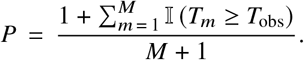

#### 4.2.5 Embedding distortion

For embedding comparisons, normalized stress was

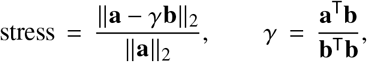

where **a** and **b** contain pairwise distances in centered CLR space and the embedding, respectively.

### 4.3. Circadian atlas

For each of the 12 tissues in GSE54650 (R. Zhang et al., 2014), we retained positive finite probes, centered their log intensities, and fit the recorded 24-h time with a first sine/cosine harmonic. The 240 probes with greatest harmonic *R*^2^, breaking ties by amplitude, were used without collapsing duplicate gene symbols. clrcycle was fit to raw positive intensities. Inferred phases were rotated and reflected against recorded time, with angular errors converted to hours. The displayed gene curves are ordinary least-squares first-harmonic fits to log intensity. Full sample feature weights are listed per tissue in Supplementary Table 1.

### 4.4. FUCCI cells and embedding benchmark

For GSE121265 (Hsiao et al., 2020), we retained the 888 cells passing the source `filter_all` field and selected the 100 genes with greatest source `trend.pve` from `fit.quant.rds.`

PCA, UMAP (McInnes et al., 2018), spectral embedding (Belkin and Niyogi, 2003), and clrcycle used the same 888×100 centered CLR matrix. PCA used two components and full SVD. UMAP used 15 neighbors, minimum distance 0.1, Euclidean distance, seed 17, and one thread. Spectral embedding used two components, a 15-nearest-neighbor graph, ARPACK, and seed 17. A circle was fit to each comparison embedding before extracting, aligning, and evaluating its phase. Stress used the original centered-CLR distances. Full feature weights per cell are listed in the Supplementary Table 2.

### 4.5. Endometrial training and held-out projection

GSE234354 counts, the seven-stage histology and 0–100 molecular time annotations of Teh et al. (2023) were mapped from Ensembl release 91 identifiers to gene symbols. The 236 annotated samples were split into 188 training and 48 held-out samples (`test_size=0.20`, seed 17), stratified by histology. Among 21,260 genes measurable in GSE234354 RNA-seq and both GSE111976 platforms, the 250 with greatest training-set variance of log(1 + *x*_*ig*_) were retained. Neither histology nor molecular time entered feature ranking or clrcycle fitting.

After fitting, the seven histological stages were mapped to midpoint days {2.5, 6, 9.5, 13.5, 17.5, 21.5, 26} and used to orient the training circle and define circular stage centers. Heldout samples were CLR-transformed, centered by the training CLR means, and projected with the fixed 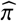
, **c**, and **s**. Stage was assigned by the nearest training center. Figure 3B colors each gene post hoc by the stage of maximum training mean centered CLR expression. Full feature weights per sample are listed in Supplementary Table 3.

#### Cross-platform projection

For GSE111976 (Wang et al., 2020), counts were summed by donor to obtain 10 10x and 19 C1 pseudobulks. Each target matrix was CLR-transformed, centered with the GSE234354 training means, and projected using the frozen gene set, 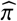, Fourier vectors, orientation, and stage centers. No target expression or annotation was used to refit the coordinates.

### 4.6. Circular feature ordering benchmark in endometrium data

We benchmarked the circular feature ordering learned by clr-cycle against a principal-circle representation and evaluated whether restricting the representation to the first Fourier harmonic discarded substantial cyclic structure in GSE234354 endometrium data. For the ordering benchmark, an ElPiGraph principal circle (Albergante et al., 2020) was fit to the centered-CLR profiles of the 250 genes across the 188 training samples, using 30 nodes, *λ* = 0.01, *µ* = 0.1, 30 iterations, 20 internal PCA dimensions, and seed 17. Genes were ordered according to their positions along the fitted principal circle, and both the ElPiGraph and clrcycle orders were aligned post hoc to each gene’s peak histological stage. Agreement with histological peak stage was quantified using circular agreement *R*, after optimal rotation and reflection. The clrcycle ordering showed greater agreement with histological peak stage than the ElPiGraph principal-circle ordering (*R* = 0.843 versus 0.770; Supplementary Fig. 2A,B).

### 4.7. First Fourier representation benchmark in endometrium data

To evaluate the first-harmonic approximation, the training expression profiles were fit as a function of cycle position using linear time, Fourier models of orders 1–3, and a periodic cubic spline with ridge regression. Model fit was quantified using in-sample *R*^2^, with no outer cross-validation, and differences in explained variance (Δ*R*^2^) were calculated relative to the first-harmonic model. The linear model explained substantially less structure than the first harmonic (Δ*R*^2^ = −0.156), whereas higher-order Fourier harmonics and the periodic spline provided only modest improvements (Δ*R*^2^ = 0.031– 0.045; Supplementary Fig. 2C,D). These benchmarks support the learned clrcycle ordering as a biologically relevant circular representation and indicate that the first Fourier harmonic retains most of the cyclic signal captured by more flexible models.

### 4.8. CHIRAL sample ordering benchmark in circadian data

We benchmarked circadian sample-phase reconstruction by clrcycle against CHIRAL (Talamanca et al., 2023) in the GSE54650 mouse tissue atlas. Both methods were evaluated using the exact tissue-specific panels of 240 probes and the same 24 samples per tissue across 12 tissues. CHIRAL was fit to log_2_ expression values with seed 17, whereas clrcycle used its native centered-CLR representation. Both inferred phase representations were aligned post hoc to collection time using the same rotation and reflection procedure. Performance was quantified using phase-resultant agreement *R* and mean absolute circular phase error in hours, with overall performance summarized by equally weighted means across tissues. Collection times informed the upstream Figure 1 probe selection and post hoc evaluation, but were not supplied during phase inference.

The two methods showed similar overall phase agreement, with mean *R* = 0.963 for clrcycle and *R* = 0.967 for CHIRAL. CHIRAL achieved a lower mean phase error overall (0.779 versus 0.862 h) and performed better on both metrics in nine tissues. In contrast, clrcycle achieved higher agreement and lower phase error in brainstem, cerebellum, and hypothalamus. In hypothalamus, clrcycle attained *R* = 0.945 and a mean phase error of 1.111 h, compared with *R* = 0.915 and 1.397 h for CHIRAL (Supplementary Table 4). This benchmark demonstrates comparable overall phase reconstruction with tissue-specific advantages for clrcycle under matched input panels and evaluation criteria.

### 4.9. Data and Code availability

Analyses were performed using clrcycle 0.1.0 https://github.com/pachterlab/clrcycle. Public dataset download scripts for all datasets used, data-processing scripts, figure generating scripts, supplementary table generating scripts and environment specifications are available at https://github.com/pachterlab/SEP_2026.

## Author contributions

N.P.S. and L.P contributed to conceptualization, performed analyses, developed the clrcycle python package, made figures, and drafted the manuscript. J.E. contributed to conceptualization and drafting the manuscript. All authors contributed to manuscript review. L.P contributed to funding acquisition.

## Acknowledgments

We thank Ali Mortazavi and Barbara Wold for valuable discussion and feedback.

## Generative AI statement

We used OpenAI Codex to assist with code refactoring and development of reproducible analysis and figure-generation scripts. OpenAI ChatGPT was also used to assist with initial drafting and language refinement of portions of the manuscript. All AI-assisted text was subsequently substantially revised and rewritten by the authors, who take full responsibility for the final content. All resulting code, analyses, figures, and manuscript text were reviewed and validated by the authors.

