## Supplementary Figures for "Dimensionality Reduction for Cyclic Compositional Data"

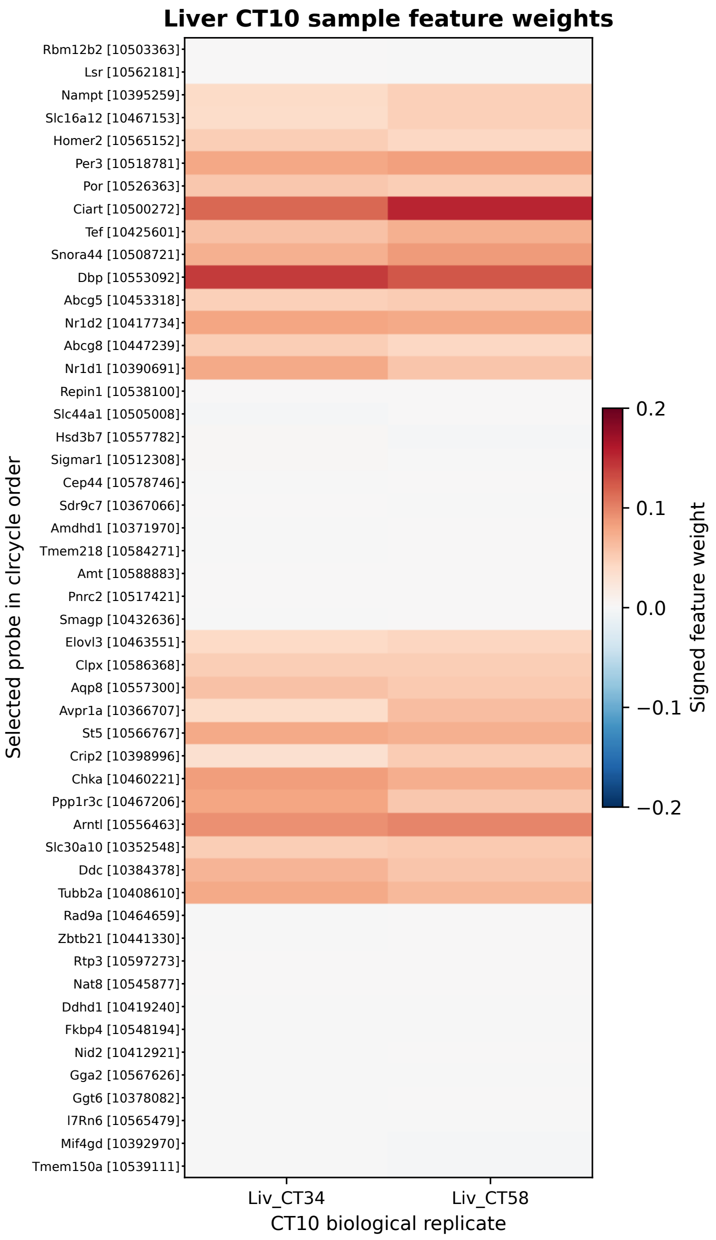


**Supplementary Figure 1: Liver CT10 sample feature weights.** Sample-specific clrcycle feature weights are shown for the two liver samples corresponding to CT10, Liv_CT34 and Liv_CT58. From the 240 probes in the fitted liver model, one probe per displayed gene was retained, choosing the probe with the largest absolute mean feature weight across the two samples. The 25 genes with the highest and 25 with the lowest mean weights were selected and displayed in clrcycle feature order without clustering. Each heatmap cell is the feature weight for one probe in one sample; replicate means were used only to select a common set of 50 probes. Positive weights reinforce the sample's projected direction, whereas negative weights oppose it. Across the complete 240-probe decomposition, feature weights sum to the sample's clrcycle radius; the displayed 50-probe subset is not expected to sum to the radius. Gene symbols and probe identifiers are shown.

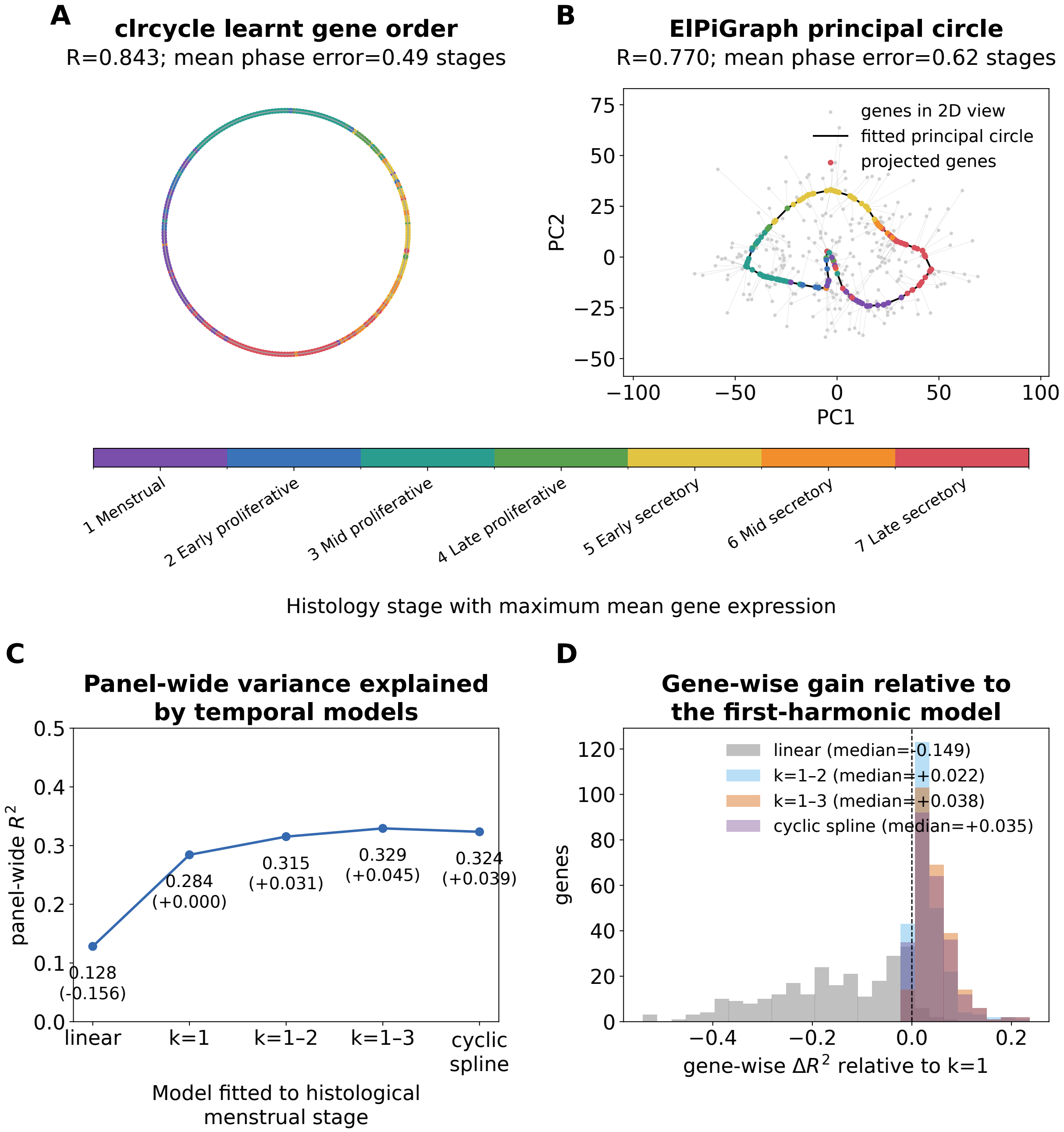


**Supplementary Figure 2: Circular-ordering and harmonic-distortion benchmarks.** **A,** clrcycle ordering of the 250 endometrial genes, colored by the histological stage with maximum mean expression. **B,** ElPiGraph principal-circle ordering of the same genes shown in a two-dimensional principal-component view. For A and B, circular agreement and mean phase error compare each ordering with histological peak stage. **C,** panel-wide variance explained by linear, first-harmonic ($k = 1$), higher-harmonic, and cyclic-spline models fitted to histological menstrual stage; values in parentheses are changes relative to the first-harmonic model. **D,** gene-wise changes in $R^2$ relative to the first-harmonic model. All methods use the same selected gene panel and samples.
